# Experimenter familiarization is a crucial prerequisite for assessing behavioral outcomes and reduces stress in mice not only under chronic pain conditions

**DOI:** 10.1101/2022.10.03.510524

**Authors:** Daniel Segelcke, Steven R. Talbot, Rupert Palme, Carmen La Porta, Esther Pogatzki-Zahn, André Bleich, Anke Tappe-Theodor

## Abstract

Rodent behavior is affected by different environmental conditions. These do not only comprise experimental and housing conditions but also familiarization with the experimenter. However, specific effects on pain-related behavior and chronic pain conditions have not been examined. Therefore, we aimed to investigate the impact of different housing conditions, inverted day-night cycles, and experimenter familiarization on male mice following peripheral neuropathy using the spared nerve injury (SNI) model. Using a multimodal approach, we evaluated evoked pain-related-, anxiety- and depression-like behavior, corticosterone metabolite levels and utilized an integrative approach for relative-severity-assessment.

Different environmental conditions are represented by individually ventilated cages and standard open cages combined with a reversed day-night-light cycle and experimenter habituation, inducing differentially modulated multidimensional pain- and emotion-like phenotypes in SNI mice. In addition, familiarization reduced the stress level caused by behavioral tests. Although no environmental condition significantly modulated the severity in SNI mice, it influenced pain-affected phenotypes and is, therefore, crucial for designing and interpreting preclinical pain studies. Moreover, environmental conditions should be considered more in the reporting guidelines, described in more detail, and discussed as a potential influence on pain phenotypes.

## INTRODUCTION

Translational pain research in animals involves several aspects within an experimental setting. Among these are the choice of the model organism, the pain model, and the measurement of clinically relevant pain modalities (outcome measures) [1,2]. Rats were the primary model species in the early days of preclinical pain research. Meanwhile, because of the availability of transgenic methods, similar numbers of mice are used, regardless of the pain models and thus the pain entities [1–4]. Mouse experiments mainly use inbred strains, such as C57BL6, which are phenotypically indistinguishable from outbred strains [5] but insufficiently reflect the heterogeneity of patients. Moreover, in half of the pain studies, mice of the male sex are currently used [1–3]. Not only the choice of animal species has changed over time, but also the pain entities that have been modeled pre-clinically [1]. Among the different existing preclinical models of neuropathic pain, currently, mononeuropathy unilateral surgical injuries of sciatic nerve branches, such as spinal nerve ligation (SNL) [6], chronic constriction injury (CCI) [7], and spared nerve injury (SNI) [8] are mainly used. These models induce localized and well-defined pain-related behaviors with a high reproducible rate for evoked pain-related behavior to mechanical stimuli across laboratories worldwide. Although all these facts represent limitations for translation, the reproducibility of studies is often limited.

While articles routinely report on the model organism, the pain model and their influence on outcome parameters, including well-being [9], the impact of the experimental environment is rarely reported and discussed. The experimental environment includes not only the behavioral test conditions (e.g., habituation period, gender of experimenter [10]) but also the animals’ housing conditions or handling methods. These factors are critical for behavioral phenotyping [11] and are hardly considered a potential source of variability in behavioral outcomes [1–3] but might affect experimental reproducibility across different laboratories.

How environmental variables might influence pain-related behavior is poorly investigated, but the modulation of the stress system is likely to be involved. Well-known examples are stress-induced hyperalgesia [11], stress-induced analgesia [12] or social transfer of pain or analgesia [13]. In addition, experimenter presence or manipulations by the experimenter usually involves an escape reaction of rodents, which is stress-associated. For example, rodents can shift from a freezing state (freeze for action) to fight-or-flight actions [14]. This escape and stress state is undesired and could affect behavioral readouts. Therefore, it should primarily be in the interest of animal welfare and the potential influence on outcome parameters to limit stress.

In particular, the choice of the cage system, the day and night light cycle and the animal familiarization with the experimenter through sufficient handling procedures might be critical influencing factors for the stress level and the well-being of the animals, and thus the severity caused by the experiment. Usually, laboratory rodents are maintained in standardized cages in designated housing rooms with particular temperature, humidity, light conditions and hygiene guidelines, which are generally determined by regulatory authorities based on guidelines and recommendations [15]. However, although rodent housing conditions are usually standardized, several differences between laboratories exist, depending on the cage system used, housing room size and capacity. Indeed, rodents can be housed in individually ventilated cages (IVC) or conventional type II polycarbonate cages, from now on, called “open-top” cages (OTC). Differences in phenotype expression between the two housing systems have already been reported and claim the need for detailed reporting of the husbandry condition [16], as required in part by the ARRIVE guidelines 2.0 [17].

The influence of environmental factors, such as ambient noise and scents, cage occupancy in the storing racks, the frequency and duration of different experimenters present in the housing room, and occupancy of the cages with other mice, especially in OTC versus IVC cages, have so far not been investigated or reported in chronic pain models. Besides the effect of the cage system, changes in the day-night light cycle are discussed to influence and potentially improve behavioral outcomes. An inverted day-night light cycle allows the study of nocturnal rodents during their active period. Considering this natural evolutionary characteristic could bridge the limitations of rodents in research and provide a better assessment of specific behavioral outcomes, such as anxiety-related behavior. However, such a refinement method has not yet been performed in preclinical pain research.

The handling method for cage changing has also been investigated and shown to affect different behavioral outcomes [18–20]. Stressful handling might contribute to the failure to replicate phenotypes within and between experiments [21], but its influence has not been studied in pain research. Since it is unclear if the environmental parameters mentioned above can represent important modifiers of pain-related behaviors, we were interested in investigating the effect of the cage system (IVC versus OTC), the day-night light cycle, and familiarization with the experimenter on pain- and emotional-related behavior in C57Bl6 mice in the SNI-model. To understand how experimenter familiarization affects stress and consequently impacts pain-related behavior, we measured the concentration of corticosterone metabolites in feces caused by experimenter handling and behavioral testing. In addition, the extent to which the environment influences the well-being and the burden of mice was investigated using the Relative Severity Assessment (RELSA) scoring system. This study is therefore essential to determine the environmental factors that need to be controlled for future study design in biomedical mouse research, particularly in pain research.

## RESULTS

### Multimodal investigation workflow

We investigated the effect of unilateral spared nerve injury (SNI) of the sciatic nerve, inducing peripheral neuropathy on the lateral aspect of the hind paw in male C57BL6/J mice (Fig. 1a) under different housing and handling conditions. SNI- and sham (control) mice were housed in two cage systems (IVC or OTC). Mice were either maintained under regular 12-hour light/ dark cycle (IVC and OTC; 7:00am light on/ 7:00pm light off) or inverted light cycle (Inv-IVC; 7:00am light off/ 7:00pm light on). Mice caged in OTC were either familiarized with the experimenter (Fam-OTC) or left non-familiarized (OTC) (Fig. 1b). To assess the pain phenotype at the multimodal level, we evaluated both stimulus-evoked nociceptive behavior (mechanical sensitivity) and gait (static and dynamic parameters), as well as the pain-associated emotional-like behavior (anxiety and depression). We first performed the elevated plus maze (EPM) test to assess anxiety-like behavior because it is one of the most sensible/delicate tests, influenced by prior experiences [22], and therefore, recommended to be performed first in longitudinal studies (Fig. 1c, d). The test was performed at d28 post-injury, where several studies report anxiety-related behavior [23]. Moreover, we measured anxiety-like behavior before the other tests because it has been reported to manifest earlier than depression-like behavior in neuropathic pain [24]. The withdrawal threshold to mechanical stimulation was determined on d34 using von Frey filaments (vF) on the lateral aspect of the affected hind paw, whereas gait analysis using the Catwalk was performed on d35 (Fig. 1c, d). Finally, depression-like behavior was measured at d38 by using the tail suspension test (TST; Fig. 1c, d).

**Figure 1:**
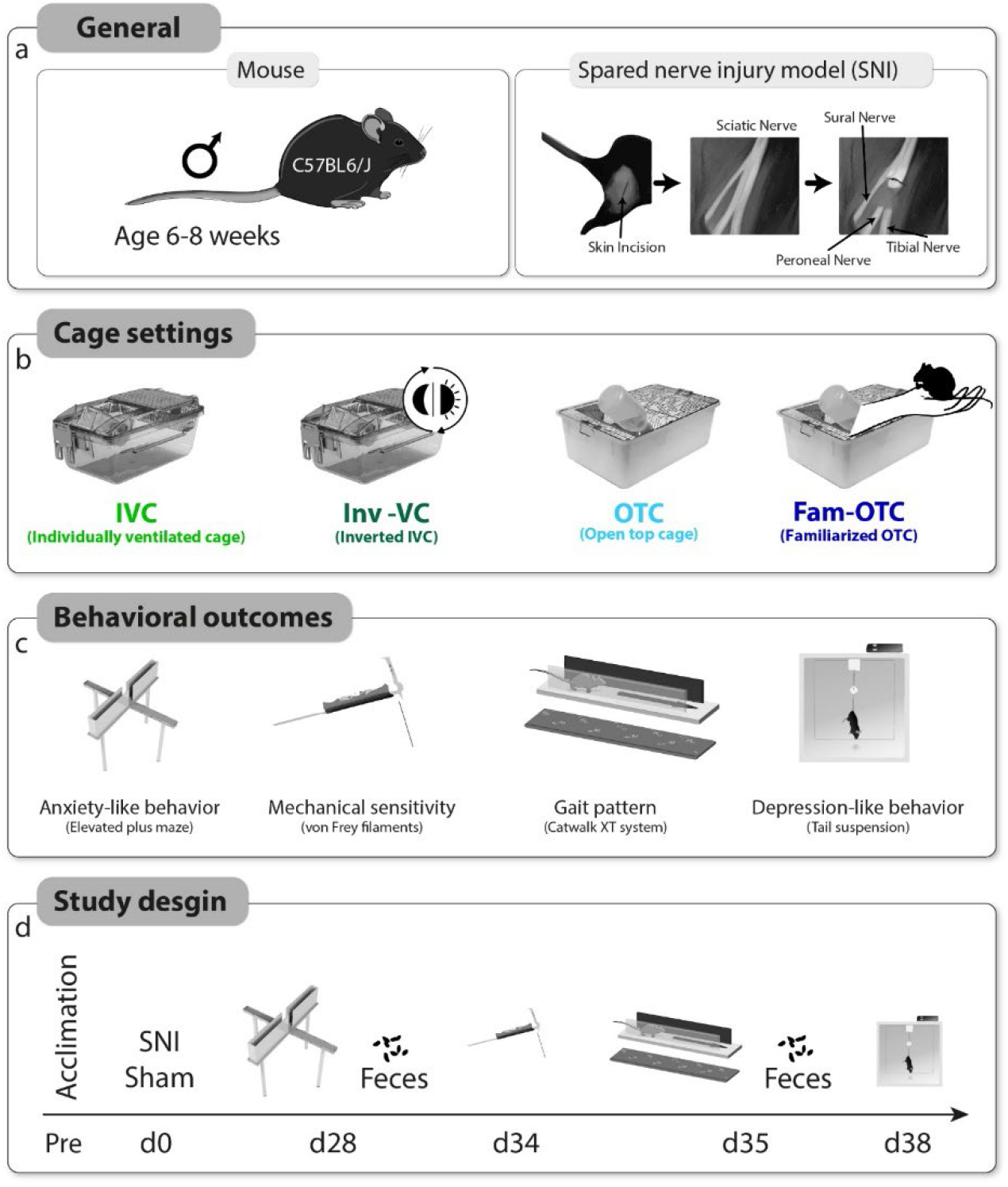
Study characteristics and design. (**a**) Male mice of strain C57/BL6/N and the spared nerve injury (SNI) model to induce peripheral neuropathic pain were used for this study. (**b**) A total of 4 different husbandry conditions were used, which differed in the inverted (Inv) of the day-night cycle in individually ventilated cages (IVC), the use of open top cages (OTC) without or with experimenter familiarization (Fam-OTC). (**c**) By using pain-, anxiety- and depression-related behavior assessments, distinct phenotypes were characterized, caused by peripheral neuropathy in combination with the four housing conditions. (**d**) An acclimatization phase was followed by induction of SNI and Sham (day (d)0). On d28, testing of anxiety behavior and collection of feces in the OTC and Fam-OTC groups was performed. Mechanical sensitivity testing was performed on d34 and gait assessment on d35 after induction of peripheral neuropathy. After gait testing, feces were collected from the OTC and Fam-OTC groups. Finally, depression-associated behavior was assessed by the tail suspension test (TST).

### Experimenter familiarization and inverted day-night cycle affect the pain-related phenotype of peripheral neuropathy

Mechanical hypersensitivity was significantly present in all SNI mice compared to the corresponding sham controls of the same experimental condition, as revealed by substantially higher response frequency to the individual vF filament stimulation and thereby significantly lower 40% response threshold in SNI versus sham mice (Fig. 2a, 2b). Interestingly, mechanical sensitivity differed between the cage systems and handling conditions and was affected in both sham and SNI mice.

**Figure 2:**
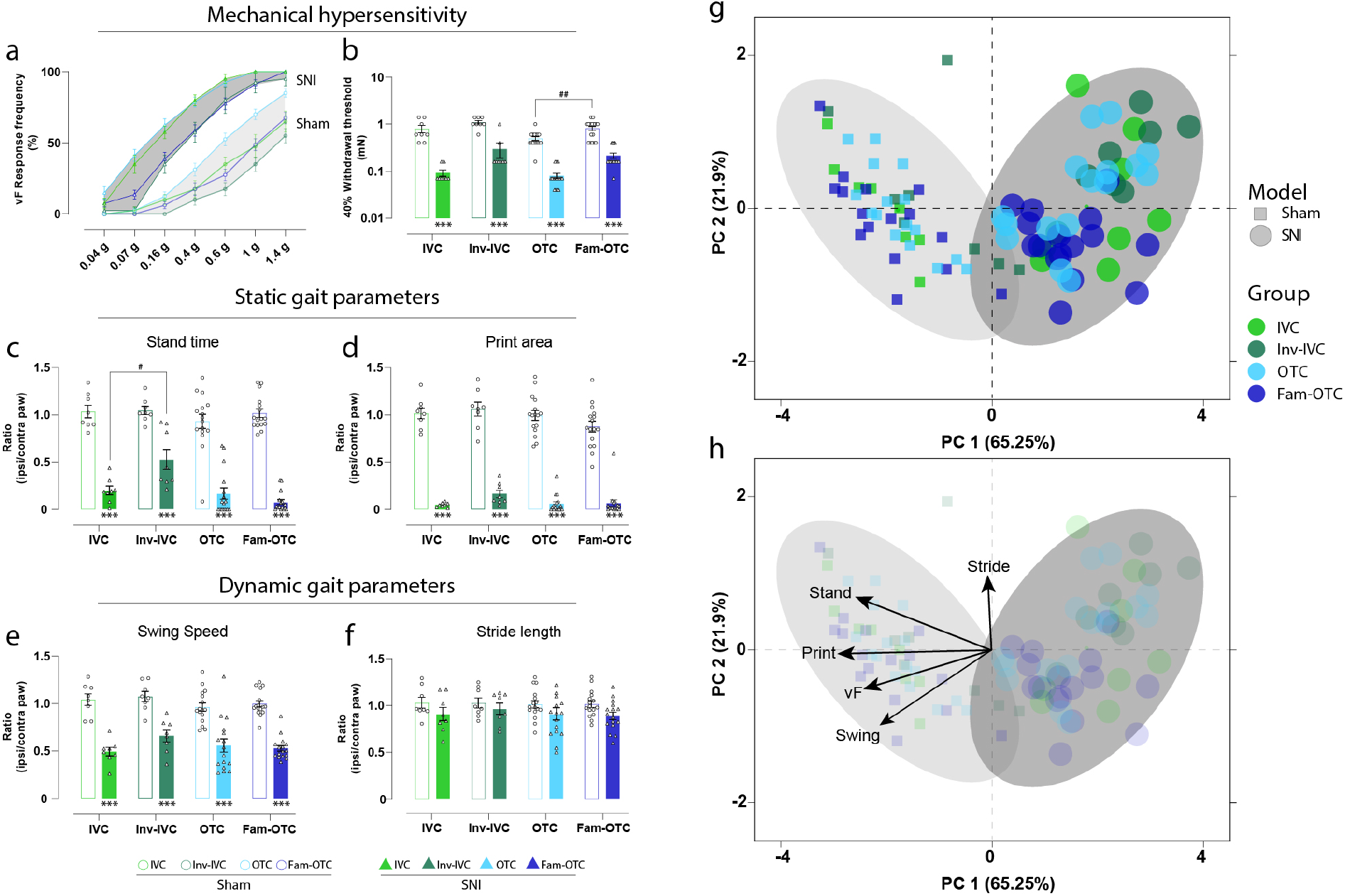
Pain-related phenotype of peripheral neuropathy is affected by experimenter familiarization and an inverted day-night cycle. (**a, b**) The response frequency in % for each vF filament and 40% response threshold in all experimental groups showed a mechanical hypersensitivity in SNI compared to sham under the different housing conditions. (**c-f**) Due to peripheral neuropathy, static and dynamic parameters were significantly modulated within the gait analysis compared to sham. An exception is the stride length, which was not changed. (**g, h**) Through the multivariate analysis of pain-associated parameters, a distinct phenotype of peripheral neuropathy (SNI) was found, which is characterized by mechanical hypersensitivity (reduction of mechanical threshold), and an antalgic gait pattern (reduction of stand time, print area, and swing time). Results are expressed as mean ± SEM. N= 8 for all IVC and Inv-IVC groups and N=16 for all OTC and Fam-OTC groups. Two-way ANOVA (repeated measures based on GLM) followed by Dunnett’s multiple comparison test. * for comparison to sham; p-values: * ≤ 0.05, ** ≤ 0.01, *** ≤ 0.001. # for comparison to husbandry conditions; p-values: # ≤ 0.05, ## ≤ 0.01, ### ≤ 0.001. The PC components were selected to determine the eigenvalues. MANOVA was used for cluster analysis in PCA (see Table 1).

Sham OTC mice showed a more pronounced mechanical sensitivity (p<0.001) and sham Inv-IVC a lower(p<0.001) mechanical sensitivity (response frequency, AUC) compared to IVC and Fam-OTC sham animals. The mechanical hypersensitivity in SNI mice in IVC was significantly enhanced compared to Inv-IVC (p<0.001) and OTC (p<0.001). Additionally, mechanical hypersensitivity was significantly higher in OTC compared to Fam-OTC (p<0.001) and Inv-IVC (p<0.001) in SNI mice (Fig. 2a). Plotting the 40% response threshold of vF-application revealed comparable results (Fig. 2b). The threshold results of the sham and SNI mice in each group were significantly changed (p<0.001) in the same direction. We observed a significant increase in response frequency (p<0.01) between OTC and Fam-OTC in the sham condition. Inv-IVC SNI mice tended to have a higher response threshold than IVC or OTC mice (Fig. 2b).

We and others have previously shown that an antalgic gait manifests in SNI mice because of unilateral peripheral neuropathy [25–28]. Therefore, we calculated and plotted the ratio of pain-related gait parameters of the affected (ipsilateral) over the uninjured (contralateral) hind paw. Regular gait pattern is reflected by a ratio of 1. We observed a significant reduction (p < 0.001) in stand time, print area, and swing speed of the affected paw in the SNI compared to sham mice (Fig. 2c-e). The stride length was not altered in any group (Fig. 2f). However, the weight bearing during gait, as indicated by the stand time of the affected hind paws in SNI mice, was significantly different (p < 0.05) between IVC and Inv-IVC housing conditions (Fig. 2c).

Next, to better understand the extent to which the cage system, the circadian cycle, and experimenter familiarization alter the multivariate pain-related phenotype of peripheral neuropathy, we applied two-dimensional principal component analysis (PCA) (Fig. 2g, h) [29]. This procedure showed that peripheral neuropathy leads to globally significant group segregation between sham and SNI independent of husbandry condition (p < 0.001; Table 1, Fig 2g). In addition, the SNI pain-related phenotype is characterized by a reduced mechanical withdrawal threshold (vF), as well as a reduction in stand Time, print Area and swing Speed (Fig. 2h). Besides global SNI phenotype, significant group segregation between IVC and Inv-IVC (p < 0.05), Inv-IVC and OTC (p < 0.05), as well as in Inv-IVC and Fam-OTC (p < 0.05) were found in SNI mice (Table 1). In sham animals, we detected significant group segregation between IVC and OTC (p < 0.05), Inv-IVC and OTC (p < 0.001), as well as in OTC and Fam-OTC (p < 0.05).

**Table 1:** Results of multivariate analysis of variance (MANOVA) in Figure 2 g, h.

| Fig. | Group | Variable | Model | Sign. | Wilks-Lambda |
| --- | --- | --- | --- | --- | --- |
| 2 | All |  | Sham – SNI | <0.001*** | 0.197 |
| 2 | IVC |  | Sham - SNI | <0.001*** | 0.102 |
| 2 | Inv- IVC |  | Sham - SNI | <0.001*** | 0.198 |
| 2 | OTC |  | Sham - SNI | <0.001*** | 0.174 |
| 2 | Fam- OTC |  | Sham - SNI | <0.001*** | 0.148 |
| 2 | IVC | Inv- IVC | SNI | 0.029* | 0.580 |
| 2 |  | OTC | SNI | 0.724 | 0.970 |
| 2 |  | Fam- OTC | SNI | 0.091 | 0.796 |
| 2 | Inv- IVC | OTC | SNI | 0.011* | 0.651 |
| 2 |  | Fam- OTC | SNI | 0.038* | 0.732 |
| 2 | OTC | Fam- OTC | SNI | 0.132 | 0.870 |
| 2 | IVC | Inv- IVC | Sham | 0.367 | 0.857 |
| 2 |  | OTC | Sham | 0.048* | 0.749 |
| 2 |  | Fam- OTC | Sham | 0.499 | 0.936 |
| 2 | Inv- IVC | OTC | Sham | <0.001*** | 0.394 |
| 2 |  | Fam- OTC | Sham | 0.147 | 0.833 |
| 2 | OTC | Fam- OTC | Sham | 0.016* | 0.752 |

### Different cage conditions produce distinct emotional phenotypes

Before applying any other test paradigm to the mice, we assessed anxiety-like behavior with the Elevated plus maze (EPM) test on day 28 post-surgery to ensure that other test manipulations did not affect the behavioral outcomes. As a result, we found a significantly increased anxiety-related phenotype in SNI compared to the sham mice only in the familiarized group (Fam-OTC), represented by a significant decrease in the open arm (OA) time (p < 0.01; Fig. 3a) and a significant increase in closed arm (CA) time (p < 0.05; Fig. 3b). No differences were observed between SNI and sham mice of the IVC, Inv-IVC or OTC groups. In addition, the inverted day-night cycle (Inv-IVC) led to a significant increase in OA time compared to the standard IVC housing conditions in the SNI group (p < 0.05; Fig. 3a).

**Fig 3:**
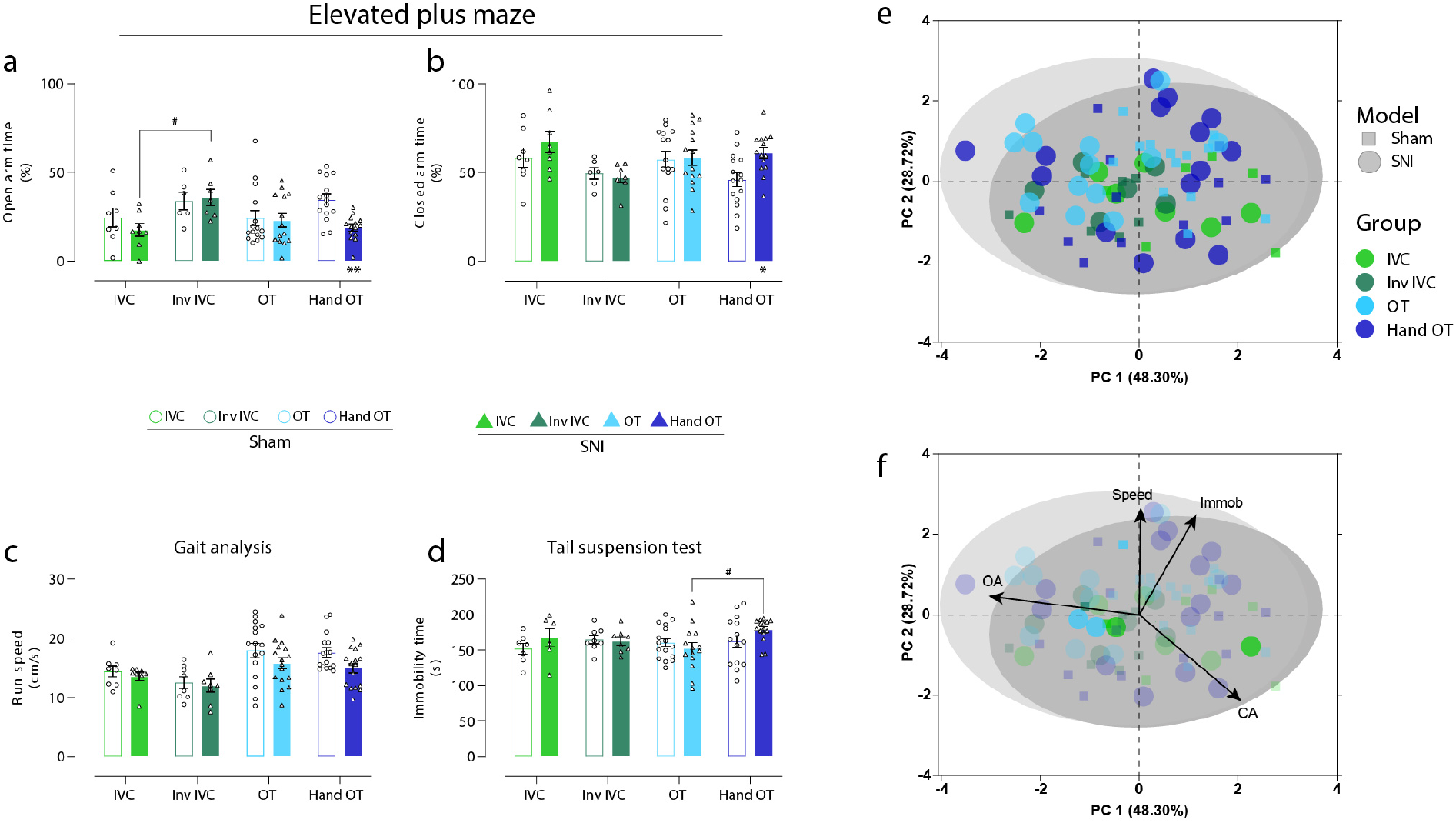
Different cage conditions produce distinct emotional phenotypes. (**a, b**) Only in Fam-IVC, a significant reduction of open arm (OA) and increased closed arm (CA) time was detected compared to sham, indicating anxiety-associated behavior. No other group showed such an effect of SNI mice compared to sham (control). A significant increase in OA time in Inv-IVC compared to IVC and a trend of reduced CA time was detected, indicating a reduction in anxiety-related behavior. (**c**) No significant change in moving speed between sham and SNI and housing conditions was observed in any group. (**d**) Depression-like behavior triggered by peripheral neuropathy was not found. However, immobility time was significantly higher in the Fam-OTC group than in the OTC group in SNI mice. (**e, f**) Multivariate analysis to characterize the emotional phenotype induced by peripheral neuropathy showed that the SNI phenotype was characterized by decreased OA, increased CA, and immobility time. Results are expressed as mean ± SEM. N = 6-8 for all IVC and Inv-IVC groups and N=14-16 for all OTC and Fam-OTC groups. Two-way ANOVA (repeated measures based on GLM) followed by Dunnett’s multiple comparison test. * for comparison to sham; p-values: * ≤ 0.05, ** ≤ 0.01, ***≤ 0.001. # for comparison to husbandry conditions; p-values: # ≤ 0.05, ## ≤ 0.01, ### ≤ 0.001. The PC components were selected to determine the eigenvalues. MANOVA was used for cluster analysis in PCA (Table 2).

Alterations in the emotional state, such as anxiety-like or stress-related behavior, could be reflected in changes in exploration behavior or running speed (Fig. 3c). To compare and correlate the general moving speed with emotional behaviors, we analyzed the running speed from the CatWalk investigation. Although we did not find significant speed differences between sham and SNI mice of all groups and between the housing and handling groups, OTC mice, irrespective of the group, displayed higher moving speed than IVC groups (Inv-IVC mice had the slowest speed). On the other hand, SNI mice of all housing groups showed a non-significant slight reduction in moving speed compared to their corresponding sham group (Fig. 3c).

As the last test, we measured depression-associated behavior on day 38 post-surgery. We could not measure any significant difference in immobility time in the tail suspension test between sham and SNI mice (Fig. 3d). However, SNI Fam-OTC mice showed significantly more immobility time than SNI OTC mice (p < 0.05; Fig. 3d), indicating a higher depression-associated behavior.

Multivariate analysis for phenotype determination based on anxiety and depression behavior showed that the SNI phenotype is characterized by reduced OA, increased CA, and immobility time (Fig. 3e,f, Table 2). In addition, besides global SNI phenotype, significant group segregation between Inv-IVC and Fam-OTC (p < 0.001) was detected in SNI mice and between IVC and Fam-OTC (p < 0.05) in sham (Table 2).

**Table 2:** Results of multivariate analysis of variance (MANOVA) in Figure 3 e, f.

| Fig. | Group | Variable | Model | Sign. | Wilks-Lambda |
| --- | --- | --- | --- | --- | --- |
| 3 | All |  | Sham – SNI | **0.041\*** | **0.919** |
| 3 | IVC |  | Sham - SNI | 0.458 | 0.855 |
| 3 | Inv- IVC |  | Sham - SNI | 0.830 | 0.963 |
| 3 | OTC |  | Sham - SNI | 0.353 | 0.917 |
| 3 | Fam- OTC |  | Sham - SNI | **0.002\*\*** | **0.606** |
| 3 | IVC | Inv- IVC | SNI | **0.037\*** | **0.517** |
| 3 |  | OTC | SNI | 0.396 | 0.884 |
| 3 |  | Fam- OTC | SNI | 0.254 | 0.843 |
| 3 | Inv- IVC | OTC | SNI | 0.376 | 0.885 |
| 3 |  | **Fam- OTC** | **SNI** | **<0.001\*\*\*** | **0.371** |
| 3 | OTC | Fam- OTC | SNI | 0.089 | 0.802 |
| 3 | IVC | Inv- IVC | Sham | 0.542 | 0.885 |
| 3 |  | OTC | Sham | 0.392 | 0.906 |
| 3 |  | **Fam- OTC** | **Sham** | **0.034\*** | **0.672** |
| 3 | Inv- IVC | OTC | Sham | 0.410 | 0.906 |
| 3 |  | Fam- OTC | Sham | 0.263 | 0.846 |
| 3 | OTC | Fam- OTC | Sham | 0.113 | 0.840 |

### Experimenter familiarization decreases fecal corticosterone metabolites

Intrigued by the effect of the experimenter familiarization on pain-, anxiety, and depression-related behavioral outcomes, we measured fecal corticosterone metabolites (FCMs) as an index of stress-hormone levels in sham and SNI mice of the OTC and Fam-OTC conditions at two timepoints; d28, reflecting the state of the stress hormone levels before any behavioral test had been performed (basal state), and at d38, reflecting the hormonal state towards the end of all experimental procedures (post procedures; Fig. 4). Interestingly, at basal d28 state, non-familiarized OTC SNI mice had significantly higher FCM values than sham mice and Fam-OTC SNI mice (p < 0.05; Fig 4a). Furthermore, following behavioral testing (post; day 38), the FCM values in non-familiarized sham mice increased significantly compared to their basal FCMs (p < 0.05; Fig. 4a). Interestingly, FCM values were not altered in Fam-OTC mice following behavioral testing and also not between SNI and sham mice (Fig. 4a). Thus, handling prevented a rise in FCM levels not only in SNI animals on their basal level; it also prevented the increase in FCM concentrations following behavioral tests in sham mice. In addition, PCA analysis revealed significant group segregation between sham and SNI under OTC husbandry (p < 0.01) and between OTC and Fam-OTC in both experimental groups (sham, p < 0.05; SNI, p < 0.05; Table 3, Fig 4b, c). These findings indicate that experimenter-dependent behavioral testing induces stress measurable via FCMs, which could be prevented by experimenter familiarization.

**Figure 4:**
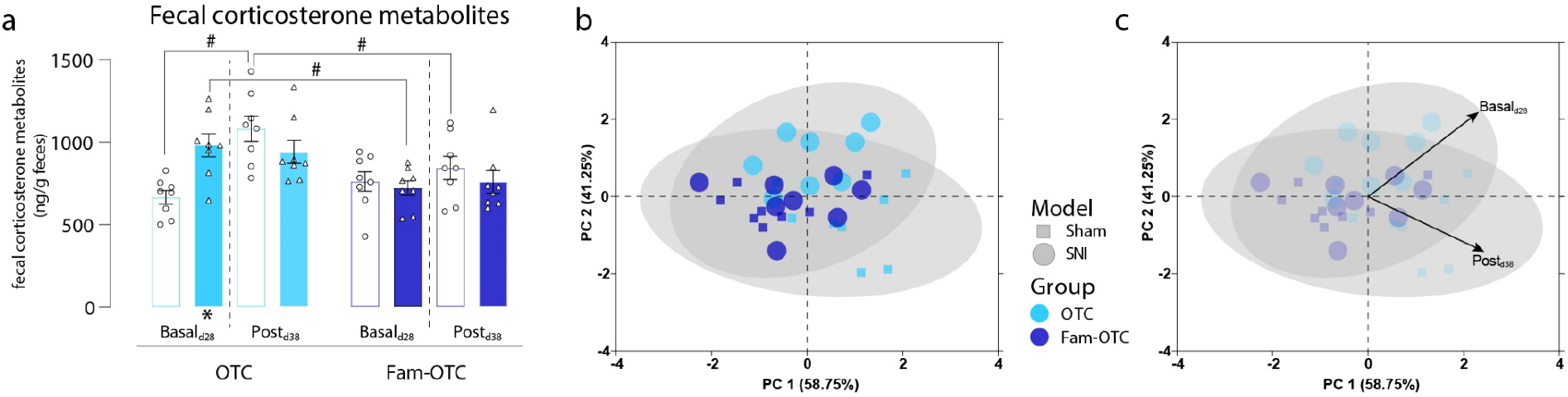
Experimenter familiarization decreases fecal corticosterone metabolites. (**a**) In SNI mice of the OTC group, a significant increase in fecal corticosterone metabolites (FCMs) was observed at d28 (basal) but not d38 (post) after the behavioral test. However, FCMs increased significantly between d28 and d38 in sham mice in the OTC group. In the group comparison of housing conditions, a significant decrease in FCM levels was detected in the Fam OTC group. (**b, c**) PCA analyses revealed significant group segregation between OTC and Fam OTC housing conditions, in both sham and SNI mice, with the OTC characterized by higher basal and post FCM values. Within the OTC group, significant segregation between sham and SNI was observed. Results are expressed as mean ± SEM. N = 8 for all groups. Two-way ANOVA (repeated measures based on GLM) followed by Dunnett’s multiple comparison test. * for comparison to sham; p-values: * ≤ 0.05, ** ≤ 0.01, ***≤ 0.001. # for comparison to husbandry conditions; p-values: # ≤ 0.05, ## ≤ 0.01, ### ≤ 0.001. The PC components were selected to determine the eigenvalues. MANOVA was used for cluster analysis in PCA (Table 3).

**Table 3:** Results of multivariate analysis of variance (MANOVA) in Figure 4 b, c.

| **Fig.** | **Variable** | **Model** | **Sign.** | **Wilks-Lambda** |
| --- | --- | --- | --- | --- |
| 4 | **OTC** | **Sham – SNI** | **0.004\*\*** | **0.431** |
| 4 | Fam- OTC | Sham - SNI | 0.709 | 0.948 |
| 4 | **OTC – Fam- OTC** | **SNI** | **0.038\*** | **0.580** |
| 4 | **OTC – Fam- OTC** | **Sham** | **0.027\*** | **0.573** |

### Peripheral neuropathy leads to increased severity revealed by the RELSA score

By acquiring and integrating nine multidimensional behavioral variables (40% <u>vF</u> withdrawal threshold, <u>Stand</u> time, <u>Print</u> area, <u>Stride</u> length, and <u>Swing</u> speed of the paw ratio during walking, running <u>Speed</u>, <u>OA</u> and <u>CA</u> time in the EPM, and <u>Immobility</u> time from the tail-suspension test) of all SNI and sham groups, an objective severity assessment score per group could be assessed (Fig. 5). For this relative severity assessment (RESLA) score, the sham-IVC group served as a reference. However, the group showed elevated (non-zero) severity because of the differences in the best-achieved values of the nine input variables within the same data (Fig. 5a). This analytical approach was chosen because no baseline information was assessed. Therefore, the severity comparisons were relative to the sham IVC reference group and not to specific time points. For example, the RELSA average of the SNI IVC mice was >1, showing larger differences in the variables than the worst measured differences in the reference group. The nature of these differences was explored by averaging the RELSA weights (RW) and visualizing them as spider plots (Fig. 5b). Thus, in the SNI IVC group, the Swing variable had the largest contribution to the RELSA (RW = 2.02), followed by Stand (RW = 1.7), Print (RW = 1.5), vF (RW = 1.31), Immob (RW = 0.81), Stride (RW = 0.79), CA (RW = 0.7), OA (RW = 0.68) and Speed (RW = 0.35). RW > 1 indicates proportionally stronger escalations in the variables than the worst case in the reference set, associated with pronounced severity in SNI mice. The SNI IVC spider plot’s total standardized area (AUC) of the RWs was 1.09. This result was more than twice the reference group’s AUC of 0.51. Moreover, the RWs in the reference group showed a different composition. Here, the Print variable showed the largest contribution (RW = 0.62), followed by Stand (RW = 0.58) and Swing (RW = 0.55). The Speed variable showed the lowest contribution (RW = 0.37), followed by CA (RW = 0.46), OA (RW = 0.48), Immob (RW = 0.48), vF (RW = 0.53) and Swing (RW = 0.55). Therefore, the difference in relative severity in the IVC group between SNI and sham was significant (p < 0.0001). This pattern remained in the other experimental groups: all sham groups were below RELSA = 1 (sham IVC AUC = 0.49; sham Inv-IVC AUC = 0.48; sham OTC AUC = 0.7; sham Fam-OTC AUC = 0.62) compared to the SNI mice, which had RELSA values > 1 (SNI IVC AUC = 1.19; SNI Inv-IVC AUC = 0.87; SNI OTV AUC = 1.05; SNI Fam-OTC AUC = 1.06). All sham vs. SNI differed significantly in the RELSA score (p < 0.001), but there was no significant difference between the sham or SNI groups of different handling or housing conditions. All IVC sham mice had lower RELSA values than OTC sham mice, and INV-IVC SNI mice had lower RELSA values than all other SNI groups. Although there were no significant differences, it can be concluded that INV-IVC conditions might reduce the relative severity assessment in neuropathic pain conditions.

**Figure 5:**
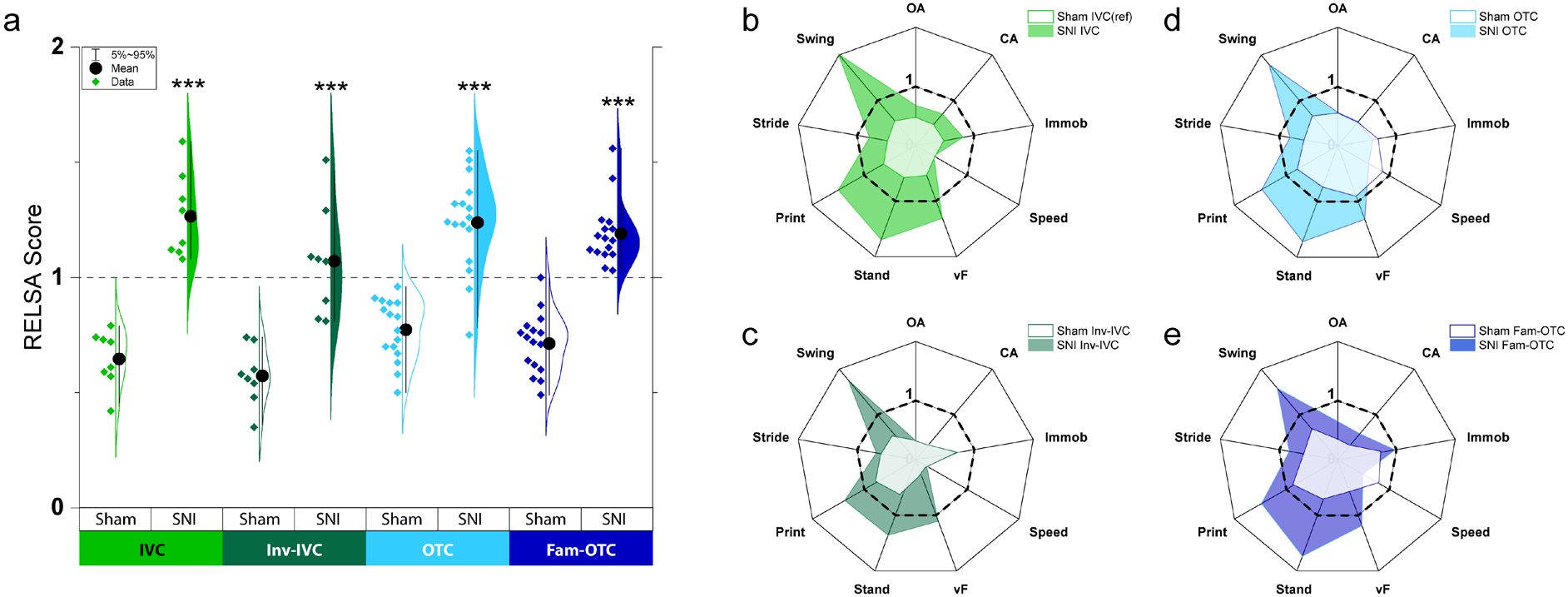
The Relative Severity Assessment (RELSA) score reveals group-dependent severity differences. (**a**) The violin plot shows the individual RELSA scores in the Sham and non-Sham groups. The average RELSA values of the Sham groups are < 1, whereas the non-Sham groups show higher severity with RELSA averages of >1. The differences between Sham and non-sham groups were significantly different, with at least p<0.05 after Tukey post hoc testing. However, no between sham or between non-Sham differences were found. The spider plots show the averaged variable contributions (RELSA weights) to the RELSA score. (**b**) In the SNI-IVC group, the Swing variable has notably the most considerable contribution to the RELSA (RW = 2.02), followed by Print (RW = 1.5), Stand (RW = 1.69), and vF (RW = 1.3). The Speed variable shows the lowest contribution (RW = 0.35). Note the consistent pattern in variable contributions in groups (**c**) to (**e**). The central areas with lighter colors show the variable contributions in the Sham groups, which are considerably smaller than the non-Sham groups (e.g., AUC_SNI-IVC_ = 1.09 vs. AUC_ref_ = 0.51).

## DISCUSSION

This is the first study comparing multidimensional pain- and emotion-related phenotypes of peripheral neuropathy in mice under different housing conditions, showing that they are modulated by familiarization with the experimenter or an inverted day-night cycle. Additionally, we could show that behavioral tests per se trigger stress and can be reduced by experimenter familiarization. Finally, the acquisition and integration of multidimensional behavioral variables revealed increased severity in SNI animals compared to sham mice by the RELSA score, which is an important tool for objectively assessing of comparable injuries in pain research and general animal research. However, the RELSA was not significantly affected by the different experimental conditions.

Pain is a multidimensional subjective experience that enables animals (and humans) to interact with the environment and its stimuli. Chronic pain conditions are associated with broader behavioral modulations, which are not limited to nociception, but also involve the emotional and affective dimension of pain and general well-being. In particular, when the pain becomes chronic the inherent warning function of acute pain is lost and maladaptive mechanisms are triggered. Changes in the patient’s emotional state reflect the maladaptive quality of chronic pain, which can manifest in complex clinical phenotypes.

To investigate the mechanisms underlying the broad clinical phenotypes of chronic pain conditions in a translational approach, multidimensional variables must be considered in preclinical studies’ design, performance and analysis.

The prerequisite and essential basis for behavioral studies is the general condition of the experimental animals. Housing conditions can have significant influences, and the experimenter-induced impact should not be underestimated [1,10]. For example, social isolation of animals via individual housing induces depression-associated phenotypes [30,31] and increases pain [32]. Conversely, social and environmental cage enrichment ameliorates chronic pain [32]. However, the effect of the housing cage type on behavior under pathological conditions, including pain, has not been studied so far. Typically, filter-lid cages connected to an automatic ventilation system (IVC) are used.

In contrast, non-filter-lid cages (OTC) are kept in standard housing racks, where the animals are more exposed to odor and noise room-environment, which might lead to deviations in various behavioral parameters between groups of animals [16,33]. Therefore, we hypothesized that environmental factors could influence mouse behavior in OTC more than in IVCs. However, we have not found a significant apparent effect between these housing conditions on pain-related behavioral outcomes.

The assessment of anxiety-related behavior is an important aspect of preclinical studies. It is controversially reported, with several factors affecting the outcome [23,34], among which an inverted day-night light cycle might be. Changing the day-night cycle and measuring behavior under darkened conditions is tricky and not often applied. Furthermore, the effect of the housing under standard or inverted day-night cycle has never been directly compared. We found a significant difference in anxiety-related behavior between IVC and Inv-IVC SNI mice but not in comparison to the corresponding sham mice. Additional factors than only inverting the light cycle might affect the phenotype. Notably, the transfer of the animals onto the maze represents a stressor per se [35].

Traditionally, handling of rats is commonly performed by hand, whereas mouse handling is mainly tail based. Different handling conditions like tail-, forceps-, cup- or tube-handling have been studied on basal behavioral parameters and aggression [18,20,36]. Tunnel or cup handling over tail or forceps handling is suggested because animals indicate less stress-induced fear and anxiety responses to human contact [18,37]. While these studies investigated the handling effect in naïve mice, the impact has not been studied under pathological conditions. Handling transiently affects corticosteroid levels and anxiety-related behavioral outcomes in an open field test conducted on the same day [38]. Handling differs from experimenter familiarization, although the terminology is not fixed. Handling means the pick-up technology from the cage, whereas experimenter familiarization is a more intense contact and getting to know the experimenter so that their interaction is not harmful. Experimenter familiarization is thought to be positive habituation of the mice to the experimenter [20,39]. Ideally, in all experimental conditions, mice should be trained to the human experimenter that their interaction is not harmful. This prerequisite is essential to ensure that acute handling stress does not affect behavioral outcomes and is supposed to increase consistency in behavioral results [40].

We hypothesized that familiarization with the experimenter would reduce stress and enable a more reliable assessment of anxiety-related and other behavioral parameters. Interestingly, experimenter familiarizing (Fam-OTC condition) enabled the measurement of anxiety-related behavior in SNI versus sham mice. Furthermore, this important finding was also reflected by depression-related behavior (measured in the TST), where Fam-OTC SNI mice displayed significantly more immobility time than OTC-SNI mice, and on the level of FCMs, which were markedly lower in FAM-OTC SNI mice and were not augmented by behavioral interference in SNI and sham mice compared to non-familiarized mice. This outcome indicates that experimenter familiarization might be the key to assessing emotional behavioral variables. Our findings are supported by a study by Ueno et al. [41], finding that repetitive experimenter handling by putting the mouse shortly on the hand of the experimenter increases the OA time of naïve C57BL/6 mice in the EPM test. Regular handling of the animals or familiarization with the experimenter is often neglected, increasing the variability and reducing the data’s reproducibility [37]. We saw an essential effect of experimenter familiarization, and a more considerable impact throughout the behavioral paradigms than the inversion of the day-night cycle, especially for emotional-like behavior testing. Therefore, we suggest experimenter familiarization as a potential prerequisite for behavioral measurements in mice.

Interestingly, both housing conditions, including day-night inversion and handling, affect the hind paws’ mechanical sensitivity in sham and SNI mice. The underlying mechanisms are likely related to stress induced by the different procedures on the animals, as we see differences in the degree and rise of FCMs between Fam-OTC and OTC mice. Overall, Inv-IVC mice, with or without peripheral neuropathy, have the highest mechanical sensitivity thresholds, followed by IVC and Fam-OTC animals, and non-handled OTC animals have the lowest mechanical sensitivity thresholds and the most pronounced antalgic gait and moving speed.

Rodent pain models are used for investigating human pain diseases and accompanied by reduced animal welfare and for ethical considerations according to the corresponding directives, need to be classified according to the severity level. Applying the RELAS enabled us to consider the severity of the peripheral neuropathy in combination with the housing conditions and the experimenter familiarization. We could show that the severity of the SNI is significantly higher compared to control mice and that the housing and experimental conditions we applied here barely influence the RELSA. Further refinement and additional modifications and the application of the RELSA in other studies will enable a better characterization of severity in rodent pain models.

## MATERIAL AND METHODS

### Animals and housing conditions

C57BL/6N male mice (Janvier, Sulzbach, Germany) were used at the age of six to eight weeks (Fig 1a). Upon delivery, mice were housed in groups of four in either Eurostandard Type II polysulfone cages (268 x 215 x 141 mm; floor area 370 mm^2^) with standard stainless-steel lids and a built-in u-shaped feed hopper (Tecniplast, Hohenpeißenberg, Germany), called open top cages (OTC) in the manuscript, or individually ventilated cages (IVC), green line GM500 (391 × 199 × 160 mm, floor area 501 cm^2^) connected to a ventilation system (Techniplast). Before neuropathic pain was induced, mice were allowed one or two weeks of acclimatization and adaptation to the new housing situation (Fig 1a). Mice had ad libitum access to food and water, under 12-hour light/dark cycle (7:00am/ 7:00pm), ambient temperature (22 ± 2 °C) and humidity (40-60 %). The general health of mice was checked daily. All procedures followed the ethical guidelines imposed by the local governing body (Regierungspräsidium Karlsruhe, Germany) and the International Association for the Study of Pain (IASP) guidelines for animal pain research. All efforts were made to minimize animal suffering and reduce the number of animals used.

### SNI model of neuropathic pain

The SNI model for neuropathic pain (Fig 1a) was induced at the right hindlimb, as described previously (Decosterd & Woolf, 2000). Briefly, under general isoflurane (in 70% O_2_ and 30% N_2_) anesthesia, the tibial and common peroneal nerves were ligated and lesioned below; the sural nerve was kept intact. Skin and muscles were opened in sham mice, but the nerves were untouched. Finally, muscle and skin were closed with resorbable surgical suture material (Marlin violet; Catgut GmbH, Markneukirchen, Germany).

### Experimental groups

We used four different housing conditions for SNI and sham mice in this study (Fig 1b):

1. IVC group: mice housed in IVC under standard light-dark cycle (light on from 07 am for 12 hours) and without handling, except for cage change and the described experimental procedures
2. Inv-IVC group: mice housed in IVC under inverted light-dark cycle (light on from 07 pm for 12 hours) and without handling, except for cage change and the described experimental procedures
3. OTC group: mice housed in OTC under standard light-dark cycle (light on from 07 am for 12 hours) and without handling, except for cage change and the described experimental procedures
4. Fam-OTC group: mice housed in OTC under standard light-dark cycle (light on from 07 am for 12 hours) and exposed to familiarization with the experimenter, which handled them twice per week, and cage change and experimental procedures. Handling started three days after delivery and caging.

Mice were allowed to recover and kept untouched except for cage changing and handling of the Fam-OTC group for four weeks. In the first experimental round, we measured eight sham and eight SNI mice in each of the four experimental groups. The same female experimenter, who also handled the mice, performed experiments. In the second round of analysis, we repeated the Fam-OTC and OTC experiments with another eight sham and eight SNI mice per group to ensure the validity and reproducibility of these data. Here, another female experimenter performed experiments and handled the animals. For the second Fam-OTC and OTC mice, feces samples were collected at two time points (d28 and d35) to analyze corticosteroid metabolites.

### Experimenter familiarization

Twice per week, with a gap of 3-4 days, mice of the Fam-OTC group were familiarized with the experimenter. This procedure started three days after delivery and caging of the mice throughout the experimental design. Therefore, the female experimenter placed her hand with gloves into the cage. Mice were allowed to explore the hand and allowed to climb into the hand. From the third handling day on, mice not going into the hand were carefully guided and placed in the hand. Handling time per cage was 20 minutes, approximately 5 minutes per mouse. For all behavioral tests, mice were carried in hand, and only for the last step (e.g., placing them into the behavioral compartments or test apparatus) they were shortly held at the tail base because it is easier to put them this way on a maze or enclosure. Mice mostly went voluntarily and directly on the hand. If not, they were briefly transferred to the hand by the tail base.

### Behavioral tests

All behavioral tests were performed between 9 am and 1 pm. Mice were brought into the behavioral room half an hour before testing began. All experimental setups were cleaned with 70% ethanol and soap if necessary and again with 70% ethanol at the end of the assessments.

### Elevated plus maze (EPM)

Anxiety-like behavior was evaluated using the EPM comprising an opaque-gray plastic apparatus with four arms (6 cm wide x 35 cm long), two open (illuminated with 100 lux) and two closed (20 lux), set in a cross from a neutral central intersection (6 cm x 6 cm) and elevated 70 cm above the ground floor (Fig 1c). The mice were placed in the maze center, and five-minute test sessions were digitally recorded. Sygnis Tracker software (Sygnis, Heidelberg, Germany) measured the time spent on open arms (OA) and closed arms (CA). Mice falling or jumping from the maze were excluded from the analysis.

### Mechanical sensitivity at the hind paws

Von Frey filaments (North Coast Medical, Gilroy, CA, USA) with increasing bending pressure forces of 0.07, 0.16, 0.4, 0.6, 1, and 1.4 g were consecutively applied to the plantar surface center of both hind paws. Mice were kept in standard plastic modular enclosures on top of a perforated metal platform (Ugo Basile, Varese, Italy), enabling the application of vF from below. Mice were acclimatized on three consecutive days for 1.5 h to the vF-compartments and 30 minutes before starting the measurements on an experimental day. Each filament was tested five times on the ipsilateral paw in the lateral aspect (innervated by the sural nerve) with a minimum 1-minute resting interval between each application, and the number of withdrawals was recorded [82,83]. Mechanical sensitivity was expressed as % response frequency to each filament or as a 40% response threshold (g), defined as the minimum pressure required for eliciting two out of five withdrawal responses.

### Gait analysis (CatWalk)

The CatWalk XT system (10.6 version) (Noldus, Wageningen, The Netherlands) was used to assess static and dynamic gait parameters. The Illuminated footprints™ technology captures the paw prints, and the CatWalk XT software calculated statistics related to print dimensions and the time and distance relationships between footfalls.

Mice were habituated to the CatWalk setup and allowed to cross the corridor during three sessions one day before the experimental measurement. Only completed trials within the defined velocity range between 10 and 20 cm/s, with a speed variance <60%, were accepted as passed runs and included in the analysis. These inclusion criteria ensured comparability across all trials. Footprints were visualized by green light emitted into the glass plate on which the rats were running, and runs were recorded by a high-speed camera (100 frames per second). Subsequently, three passed runs for each mouse were semi-automatically analyzed for two selected static (stand time and print area) and dynamic (swing speed and stride length) gait parameters. Additionally, running speed in cm/s was sampled for each run.

1. Print area: area of the whole paw
2. Stand duration (s): duration of ground contact for a single paw.
3. Stride length (cm): distance between successive placements of the same paw.
4. Swing speed (cm/s): rate at which a paw is not in contact with the glass plate.

### Tail suspension test (TST)

Depressive-like behavior was evaluated with the TST. Mice were suspended by the end of their tail with adhesive tape 50 cm above the surface. An approx. 1 cm thick metal rod attached to a laboratory stand with a tripod clamp served as a suspension base. The distance to the frame was 40 cm, and mice were video-monitored. As previously described, the immobility time was manually measured during the whole testing period of 6 min [16]. Mice climbing up their tail were excluded from the experiment.

### Fecal corticosterone metabolite (FCM) measurement

The measurement of FCMs is a non-invasive method to address rodent hormonal stress levels [42]. Feces were collected around noon at two time points: directly after the EPM test (day 28; reflecting basal level) and directly after the Catwalk test (day 35; reflecting post-behavioral state). Since fecal FCM peak values are reached 4-10 hours after interference [43], the direct effect of the test performed just before feces collection on FCM is excluded. Hence, the FCM levels on each collection day reflect only the cumulative effect of the different housing conditions, handling, surgery, and experimental procedures performed before. To collect a minimum of 5-6 fecal boli, mice were placed individually in empty Macrolon Type II cages (Tecniplast) for max. 60 min. According to Palme et al., the feces were stored in polypropylene tubes at −20 °C until extraction [44]. Briefly, fecal samples were dried for two hours at 80 °C before mechanical homogenization, and 0.05 g were extracted with 1 ml 80% Methanol for 30 min on a vortex. After centrifugation for 10 min at 2500 *g*, 0.5 ml supernatant was frozen until analysis. FCM was measured using a 5α-pregnane-3β,11β,21-triol-20-one enzyme immunoassay (EIA) [43].

### Statistical analyses

Statistical analyses were performed with Graph Pad Prism. The differences were considered statistically significant when P< 0.05.

For all behavior parameters, including RELSA datasets, two-way ANOVA with Dunnett’s posthoc test was used to compare groups (to sham) and for between-group analyses.

Multivariate behavioral data were analyzed by principal component analysis (PCA) with prior standardizations. Principle component (PC) selection was based on the largest eigenvalues. The first two PC were plotted as biplots. Groups were added to the biplots for illustration but were not used during the PCA. The significance of group segregation was determined by multivariate analysis of PC loadings regarding the group with Tukey post hoc tests. Multivariate ANOVA (MANOVA) was performed for further analysis.

The Relative Severity Assessment (RELSA) score was calculated with the RELSA package in the R software (v4.0.3 [45,46]). The Sham-IVC data were chosen as reference data. The current data did not include time or baseline information. Therefore, all data were standardized to the best-achieved results in the reference data, recognizing the direction of severity in each of the nine input variables (OA:max, CA:min, Immobility:min, Speed:min, vF:max, Stand:max, Print:max, Stride:max, Swing:min indicate the best-achieved results). This approach resulted in [100%; 0] scaled data, in which 100% showed no difference to the best case in the reference data. Subsequently, the experimental groups were mapped against the reference set using the RELSA algorithm, resulting in individual scores that could also contrast the experimental groups’ relative severity. For example, a RELSA score of 1 represents the case in which all nine input variables would have reached the maximum severity (worst-achieved results) in the reference set. RELSA scores >1 indicate animals with higher severity than the reference set. The individual variable contributions to the RELSA score (RELSA weights) were used to construct spider plots. In this representation, the weights were averaged per variable in each experimental group. A significance level of p < 0.05 was considered. Data were analyzed by Prism software, version 8. (GraphPad, USA), SPSS22 (IBM, USA) and OriginPro 2022 SR1 (OriginLab, Northampton, Massachusetts, USA.).

### Data availability

Primary data and datasets generated and/or analyzed during the current study are available from the corresponding author upon reasonable request.

## ACKNOWLEDGMENTS

The authors thank Julia Speck, Florencia Garrido Charrad and Edith Klobetz-Rassam for their excellent technical help. The work was performed at the Interdisciplinary Neurobehavioral Core (INBC), University of Heidelberg. This work was funded by a Collaborative Research Center 1158 (SFB1158) grant from the Deutsche Forschungsgemeinschaft (DFG) to ATT (project S01).

## AUTHOR CONTRIBUTION

DS analyzed data, performed PCA, generated figures and drafted the manuscript, SRT performed RELSA, discussed data and drafted the manuscript, RP performed FCMs analysis, CLP helped conceptualize the study and critically examined data, and, EPZ and AB critically discussed data, ATT conceived and designed the study, received funding, performed experiments, analyzed data and wrote the manuscript.

